# On-Farm Genetic Diversity of Wheat (*Triticum aestivum spp*.) In Digalu Tijo District, Arsi Zone, Ethiopia

**DOI:** 10.1101/2022.03.07.483220

**Authors:** Fekadu Gelelcha, Fikadu Kumsa, Tolera Kuma

## Abstract

**Introductory:** This study was undertaken to assess the on-farm genetic diversity of wheat Landraces (*Triticum aestivum spp*) in Digalu and Tijo District.

**Methods:** For this study, three farmer households kebeles were selected by cluster sampling (area sampling) method because they are found in the most dominant wheat landraces growing agro-ecological areas. Data for this study was collected using questionnaires, field observation, interviews, focus group discussion, and document analysis.

**Results and Discussions:** In total, 27 wheat landraces were reported to be grown before two decades to the present day. Of these, only 9 of wheat landraces are still cultivated by farmers. Among these, ‘Qamadi guracha’ is the most common growing landraces in the study area. The Shannon diversity index (H’) of growing wheat landraces ranges from 1.09 to 1.37 among the groups. The estimated genetic erosion for wheat landraces was found to be 66.7% due to the major factors: improved wheat varieties; the introduction of other more productive crops and wheat landraces low productivity.

**Conclusions:** Food quality, Pests (example, birds), diseases resistance, market value, and straw quality were factors that initiated the farmers to maintain the genetic diversity of landraces on their farmlands. However, the preservation of wheat landraces are influenced by bottlenecks like the seed selection system, and insufficient crop yield.

**Recommendation:** Regeneration of soil fertility, restoring of lost landraces, improvement of landraces, on-farm conservation by re-sowing, saving of seeds for future, and *ex-situ* conservation is suggested for the restoration of wheat landraces diversity in Digalu and Tijo District.

## 1. Introduction

Genetic diversity is usually thought of as the amount overall of genetic variability (Brown, 1983). On-farm genetic diversity also refers to the relative distribution of plant species (Adugna Abdi and Zemede Asfaw, 2005). Crop variety, in turn, refers to variation within the same and between different crops that include staple crops such as wheat in Ethiopia (Kumar *et al*., 2011; Tillman, 2001). Wheat (*Triticum aestivum*) is the grains and the first crops to be domesticated and cultivated by human beings. Most wheat varieties in Ethiopia are landraces (Efrem Bechere *et al*., 2000; Firdissa Eticha *et al*., 2006) and are traditionally grown (Endashew Bekele, 1984).

Wheat varieties whose morphological and genetic composition is shaped by household farmers’ practices, natural selection pressure over generations of cultivation, and human selection (Smale, 2001). They display genetic variation for useful quantitative and qualitative characters (Brown, 1983; Harlan, 1992, Porceddu *et al*., 1988). Even if the wheat landraces are being genetically eroded, they are produced at altitudes of 1800–3500 m.a.s.l in Ethiopia (Tesfaye Tesemma and Getachew Belay, 1991). The genetic erosion of tetraploid is the loss of wheat landraces from areas they adapted due to the introduction of productive semi-dwarf cultivars, improved varieties, and high selection pressure applied in breeding programmers (Skovmand *et al*., 2005; Royo *et al*., 2009). On-farm genetic conservation is the continued cultivation of landraces genetic diversity by household farmers in centers of domestication allows them to adapt to continually changing environmental conditions (Adugna Abdi and Zemede Asfaw, 2005). Even today, it is evident that Ethiopian farmers are practicing traditional farming of wheat landraces (Smale, 2006).

Moreover, knowledge about the level and extent of genetic diversity in wheat landraces on the farm is of great value for sustainable maintenance and utilization of genetic materials (Fassil Kebebew *et al*., 2001; Teklu Tesfaye and Hammer, 2006). Estimating any possible loss of genetic diversity and its causes; tracing the available genetic variability is important for *in situ* or *ex situ* conservation of wheat landraces (Berg and Efrem Bechere, 2007). Thus, the documentation, estimating genetic erosion and its causes, and incentives why farmers conserving landraces were the motives that initiated the present study to assess the on-farm genetic diversity of wheat *landraces* (*Triticum aestivum spp*.) for successful conservation in *in-situ* and *ex-situ* and sustainable utilization of the genetic materials of varieties in Digalu Tijo District, Arsi zone, Ethiopia. The objective of the study was to assess the on-farm genetic diversity of wheat landraces (*Triticum aestivum spp*.) in Digalu Tijo District, Arsi Zone, Ethiopia.

## 2. Material and Methods

### 2.1. Description of the study area

Digalu Tijo is one of the 25 districts found in the Arsi zone of Oromia Regional State. Its borders are Tiyo to the north, Tena to the east and Munessa to the west, and Lemmu-Bilbilo woreda to the south. It is about 198 km south of Addis Ababa and 98km from Adama town. The district contains one urban town and 27 rural kebeles i.e. 28 kebeles in total within a 92,698.51 km^2^ area. It is located at the latitude of 7°35’57” - 7°55’43” N and longitude of 38°59’40” - 39°24’31” E (Fig. 1); and its elevation ranges from 2000 - 4000 m.a.s.l. The District’s *woynadega* agroecology kebele represents areas with mid-altitude (2000 - 3000 m.a.s.l.), and it receives annual rainfall that ranges 900-2000 mm and its mean annual temperature ranges from 18-25 °C. The *dega* agroecology kebeles represent areas with altitude (3000 to 3500 m.a.s.l) and they are receiving mean annual rainfall and temperature ranges from 1500-2600mm and 10.9°C - 21 °C (Ministry of Agriculture, 2007; Tadele Ferede *et al*., 2010). 2007, the national census reported that the total population for Digalu and Tijo district was 140,466, of whom 69,503 were males 70,963 were females, and 14,080 or 10.02% of its population were urban dwellers.

**Fig.1.**
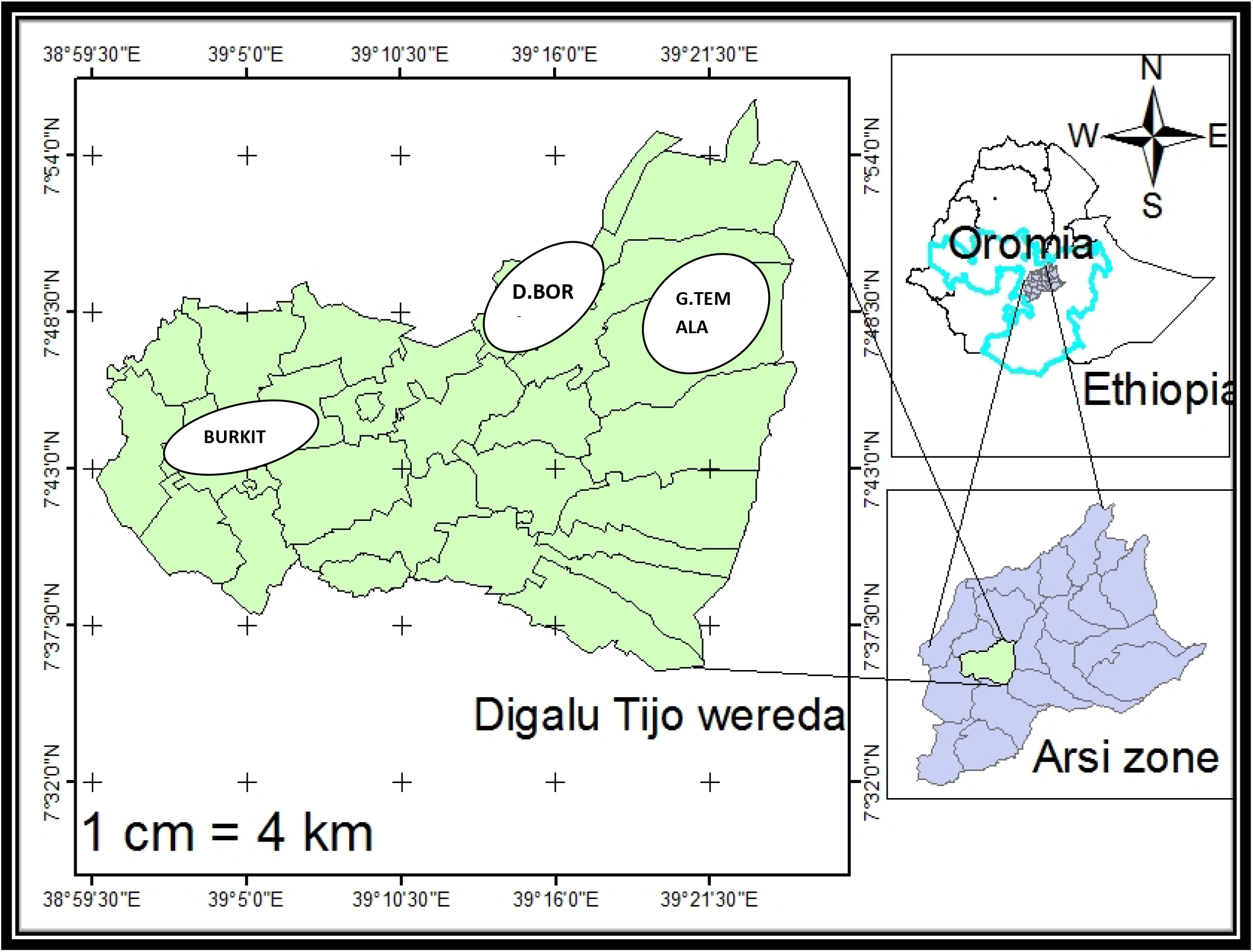
The map of Digalu and Tijo District.

#### 3.1.1. Geology and Soil type in the study area

According to MoWR (2002), the geomorphology of Ethiopia is variable and it is generally the result of repeated volcanic and tectonic events with the associated erosion of Mesozoic sedimentary and volcanic rocks, and deposition process. According to DigaluTijo district Agricultural Office Annual Report (2015), its physiographic diversity is characterized by highlands and flat-topped plateaus. Regarding its soil types, about 44% of soil type is red, 35% of soil type is black and 21% is brown soil types DigaluTijo district Agricultural Office Annual Report (2015).

### 3.2. Study design

This study was conducted from September to November 2016 because these months were the cropping seasons when the wheat landraces become well-grown and easy to get the real sample in the study area. Owing to this, the researcher selected these months to collect a reliable sample of wheat landraces and data. A descriptive survey research method was employed because it is suitable for describing the existing situation and investigating phenomena in their natural setting and it presents opportunities to combine both quantitative and qualitative i.e., to assess the on-farm genetic diversity of wheat landraces cultivated by farmers. To do this, two *kebeles* from *Dega* and one *kebele* from *woynadega* were selected by using the purposive sampling technique because they are found in the dominant wheat-growing agroecology area.

### 3.3. Study subject /target groups

The source of the population for the study was household farmers in the DigaluTijo district of three sampled kebeles. This was because farmers are found in the kebeles of dominant wheat landraces growing agroecology areas. In this case, both the male and female household farmers were considered as some female household farmers present in these kebeles.

### 3.4. Sample size and sampling methods

There were 4832 identified total household farmers in the three sampled kebeles of the study area (CSA, 2007).

Among these, the sample size (n) determination for the study was carried out through the following formula cited from Kothari (2004) as follows:

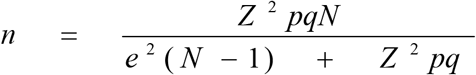

Where, **n** = the desired sample size;

**N=** the sizethe of household population in the three sampled kebeles (4832) at a Confidence level of 95% and 5% precision,
**Z =** the critical value containing the area under the normal curve =1.96
**e** = the desired precision level (5% precision = 0.05)
**p** = an estimated proportion attribute present in the population (0.1) and
**q** = 1-p (1 – 0.1 = 0.9)

By substituting these values in the above formula the sample size ‘n’ was calculated as follows:

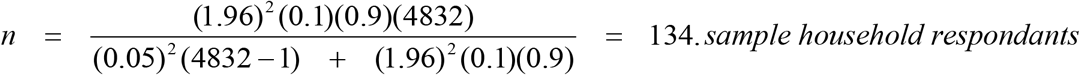

Thus, a total of 134 household farmers were determined as sample size. In addition, 9 DA workers and 3 district agricultural experts were selected. A stratified proportional probability sampling technique was used to divide and allocate the sample size (134) to each of the three selected sample kebeles as it did not constitute a homogeneous group.

Accordingly, the principle of probability is proportional to size, which means kebeles that, had large numbers of farmer households, based on the measure of size, were given greater probabilities. Then, the proportional allocation formula was employed by dividing the number of household farmers from each kebele/ stratum to the total study population and multiplying by **n**.

The formula for proportional allocation from the stratum/kebele = **n*P_i_**

Where **n**= the sample size selected from three sample kebeles in this case 134 head of households, **P_i_**= the proportion of household population includes from the stratum/kebele. Based on this, Gushatemala (**n**_= 1371_ households) = 134* 1371/ 4832 = 38 sample respondents, Burkitu (**n** _=1852_ households) =134 *1852/4832= 51 sample respondents and Digalu bora (**n** _=1609_ households) =134*1609/4852 = 45 sample respondents. The result was summarized in (Table 1).

**Table 1.** Total number of household farmers and selected sample size from each three kebele

| Sampled kebeles | Total households farmers (N) | Respondent households(n) |
| --- | --- | --- |
| Gushatemala | 1,371 | 38 |
| Burkitu | 1,852 | 51 |
| Digalubora | 1,609 | 45 |
| **Total** | **4,832** | **134** |

### 3.5. Sources of data and instruments of data collection

#### 3.5.1. Sources of data

Data were collected from both primary and secondary sources. The primary data were collected from household farmers; DA workers and Woreda Agricultural Experts. To enrich the information gathered through other tools, secondary data were collected from previously published research papers, Arsi/ Kulumsa/ Agricultural Research Institute, websites, reports, journals, books, and other thesis results.

#### 3.5.2. Instruments of data collection

To gather reliable, rich, and deep data from the subjects of study on on-farm genetic diversity of wheat landraces, the researcher employed the following five data collecting instruments:-

##### 3.5.2.1. Observation Checklist

To assess critically about on-farm diversity of wheat landraces at the time of the study, which isto known, be highly cultivated, field observation was employed in the study area.

##### 3.5.2.2. Questionnaire

To collect data from134 sampled farmers, the semi-structured questionnaires with open-ended and closed-ended questions were prepared and distributed. During data collection, a questionnaire was translated to Afaan Oromo.

##### 3.5.2.3. Interview

To acquire detailed information and to fill the gap that was not be covered by the questionnaire, an individual interview was used. In this case, the eight open-end of interviews preferred to collect data from DA and the district’s Agriculturally Experts. During data collection, interview guideline was used; and it was translated to Afaan Oromo.

##### 3.5.2.4. Focus Group Discussion (FGD)

A total of three group discussions were conducted (one in each kebele) with ten key informants to reinforce the questionnaires and find out any hidden information that could be missed. To do this, farmers were assigned based on their farming experiences of wheat landraces and made them discuss and investigate the main issues of associated questions.

##### 3.5.2.5. Documents Analysis

To enrich the information gathered through other tools; and to discover the existed facts about wheat landraces in the study area, document analysis was conducted. The researcher prepared a checklist as a guideline to conduct the analysis data obtained from documents.

### 3.6. Data management and analysis procedure

The data gathered through all the data gathering tools was analyzed with quantitative and qualitative data analysis method (numerically and narrative) systems. Following this, data gathered through document analysis and FGD were analyzed based on the specific categories set concerning the research questions.

Quantitative data were organized according to their sequences-coded, tabulated, and analyzed using statistics software (for example, correlation bivariate and descriptive statistics). The qualitative data was organized and triangulated by cross-checking from all tools and evaluating the results of the quantitative findings in different ways.

#### 3.6.1. Analysis of genetic diversity of wheat landraces

The data gathered through all the data gathering tools about the diversity of wheat landraces was analyzed systematically with quantitative (presented in numbers) and qualitative (described based on some quality) data analysis method systems. In addition,

The Shannon diversity index (*H’*), is a useful measurement of diversity, and has better discriminant ability in the situation when the number of varieties and their proportional abundance remain constant (Stiling, 2002), estimated as follows:

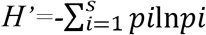

Where,

S = total number of species

Pi = is the proportion of each Variety in the sample

*l*n = log base n

Obtained results were compared diversity indices, i.e., (low, H’= 1.1, medium, H’= 3.5 and high, H’= 4.5) (Kent and Coker, 1992).

#### 3.6.2. Sorensen similarity index

A similarity that relies on the presence or absence of the wheat landraces among sampled kebeles was measured by the Sorensen similarity index which is the most common binary similarity coefficient. The obtained results were compared with the Sorensen similarity index that should not be greater than 0.5 (Kent and Coker, 1992).

Sorensen’s coefficient is expressed as **Ss = 2a/(2a+b+c)** Where

**a** = number of landraces common to the two kebeles
**b** = number of landraces unique to the first kebele
**c** = number of landraces unique to the second kebele

Often, the coefficient is multiplied by 100 to give a percentage similarity index.

Dissimilarity is then computed as: **Ds = b + c/(2a+b+c) or 1 – Ss**

#### 3.6.3. Correlation analysis of agronomic features of wheat landraces

To examine the presence of an association between the number of wheat landraces and farmlands size, correlation analysis was computed. According to (Zawdie Bishaw *et al*. 2014), a positive association is expected between large-sized farmlands and the number of wheat landraces because on wider farmlands farmers can produce more wheat landraces. Additionally, the presence of association among maturity time and productivity in kg/ ha, average stem length and productivity in kg/ha, and maturity time and average stem length of wheat landraces in the study area, Pearson correlation relationship were computed using SPSS, version 16.The results were compared with standard ‘r’ values, (r≥ 0.9= very strong relationship, 0.8 ≤ r<0.9 = strong relationship and 0.7 ≤ r< 0.8 = acceptable relationship (Kothari, 2004).

#### 3.6.4. Estimation extent of on-farm genetic erosion

The present extent of on-farm genetic erosion of wheat landraces in the study area was calculated using the formula of Hammer *et al*. (1996) and it is given as:

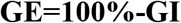

Where **GE** is genetic erosion and **GI** is genetic integrity which is given as:

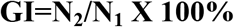

Where **N_1_** is the number of landrace varieties collected in previous times and **N_2_** is the number of presently collected landrace varieties.

#### 3.6.5. Preference ranking

Preference ranking was made for 9 wheat landraces in terms of their end-use purposes by 9 key informants. Each rank was given an integer value1-9 and the most important item was given the highest value (9), while the least important assigned the smallest value (1). The informants were given a list of wheat landraces and asked to rank them from the highest to lowest (least) in decreasing order. The rank of each variety was determined by adding up these values for all informants.

## 3. Results

### 4.1. Demographic characteristics of the respondents

Demographic Characteristics of the 134 household farmers were collected in the study area. This was based on respondents’ sex, age, educational background, family size, and farming experience in years (Table2).

**Table 2.** Demographic characteristics of the farmers

| Items | Alternatives | Respondents | Percentage |
| --- | --- | --- | --- |
| Sex | Male | 120 | 89.5% |
|  | Female | 14 | 10.5% |
|  | **Total** | **134** | **100%** |
| Age in year | 25-35 | 20 | 14.9% |
|  | 36-45 | 72 | 53.7% |
|  | 46-55 | 26 | 19.4% |
|  | 56 and above | 16 | 11.9% |
|  | **Total** | **134** | **100%** |
| Educational Background | Illiterate | 0 | 0%% |
|  | Read/Write | 71 | 52.98% |
|  | Primary | 58 | 43.3% |
|  | Secondary | 4 | 0.03% |
|  | Preparatory | 1 | 0.007% |
|  | College | 0 | 0% |
|  | University | 0 | 0% |
|  | **Total** | **134** | **100%** |
| Family size | 2-3 | 13 | 9.7% |
|  | 4-6 | 80 | 59.7% |
|  | 7 and above | 41 | 30.8% |
|  | **Total** | **134** | **100%** |
| Farming experience in year | 1-6 | 10 | 7.5% |
|  | 7-12 | 37 | 27.6% |
|  | 13 and above | 87 | 64.9% |
|  | **Total** | **134** | **100%** |

### 4.2. Wheat varieties cultivated by farmers

#### 4.2.1. Modern wheat varieties

A total of 34 wheat varieties were documented. Among these, only 7(20.7%) types of improved wheat varieties were grown mainly in the study area presently. These were described by farmers based on their phenotypic traits (example, seed color, and plant height), maturity time, end-uses quality, and yield in quintal per hectare (Table 3).

**Table 3.**
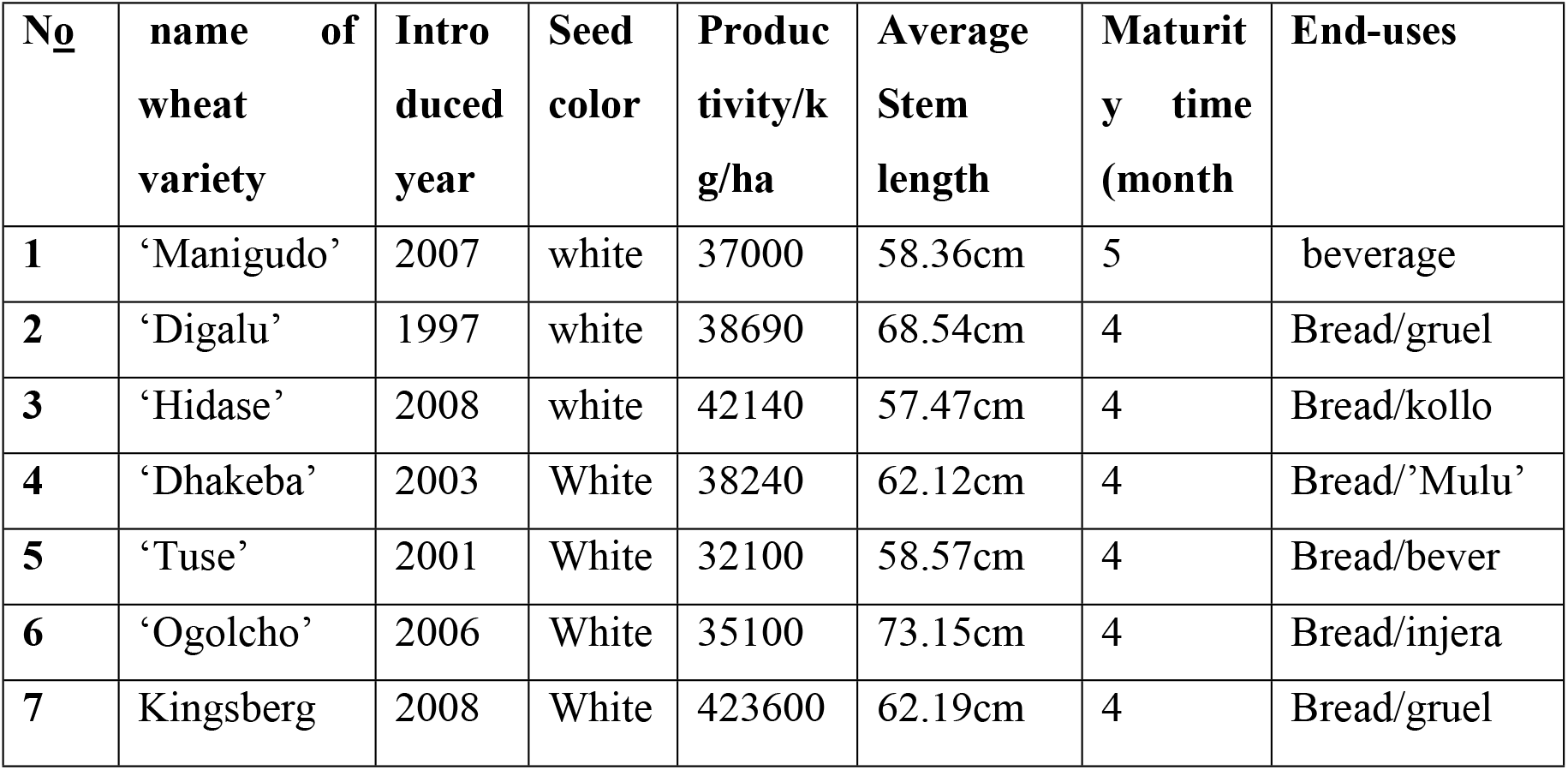
Improved wheat varieties and agronomic features in the study area

### 4.3. On-farm genetic diversity of wheat landraces

#### 4.3.1. Genetic diversity of wheat landraces

A total of 27 wheat landraces were identified by farmers of the three sampled kebeles of the study area. From those identified landraces, 18(66.6%) of them were lost and only 9(33.3%) are being cultivated presently by farmers

#### 4.3.2. Distribution of existing wheat landraces

Of currently being cultivated wheat landraces in the study area, ‘Qamadiguracha’ is the most frequently being cultivated and common to three sampled kebeles; whereas ‘Ganabasi’ was the least and rarely being cultivated (Fig. 2)

**Fig.2.**
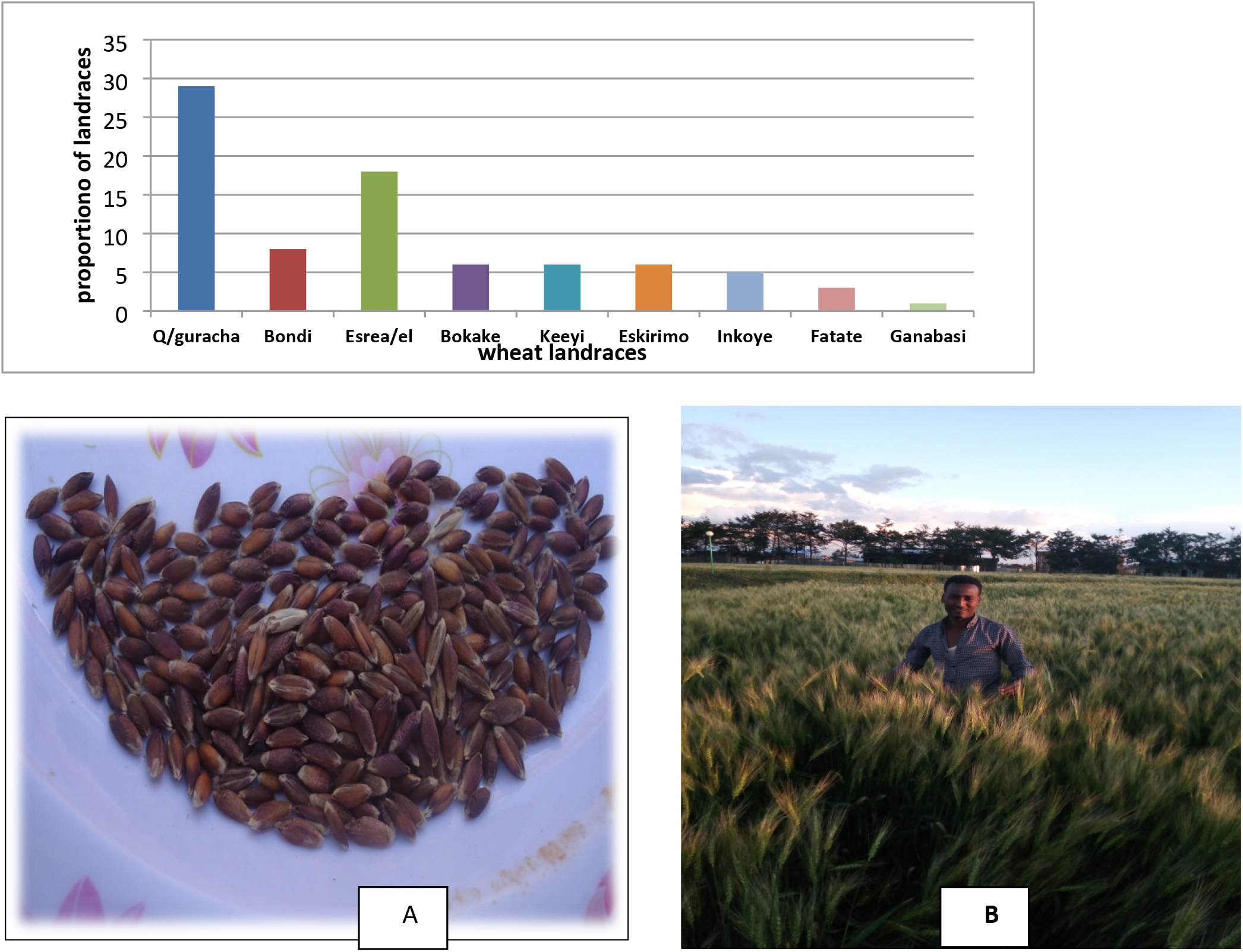
’A’ Black -seeds and ‘B’ straw of ‘Qamadi guracha’

**Fig.3.**
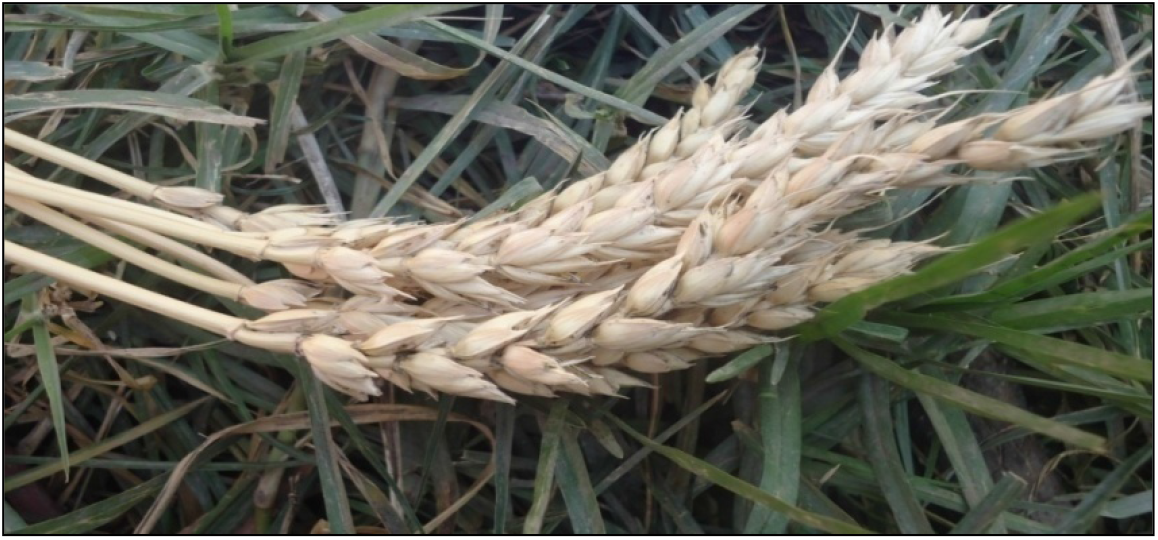
The non-spikes, ‘Bondi’

#### 4.3.3. Analysis of wheat landraces diversity by Shannon index

The diversity of wheat landraces was estimated based on the number of varieties collected. As a result, the Gushatemala kebele showed the highest diversity (H’= 1.37) followed by Digalubora kebele (H’= 1.32). Burkitu kebele was found to be less diverse in terms of the number of varieties collected in this study area (H’= 1.09) (Table 4).

**Table 4.** List of wheat landraces collected along with diversity estimate.

| kebele | Burkitu |  | Digalubora |  | Gushatemala |  |
| --- | --- | --- | --- | --- | --- | --- |
| indices |  |  |  |  |  |  |
|  | Wheat landraces | Frequency | wheat landraces | frequency | wheat landraces | Frequency |
|  | ‘Qamadiguracha’ | 5 | ‘Qamadiguracha’ | 12 | ‘Qamadiguracha’ | 17 |
|  | ‘Bokake’ | 6 | ‘Inkoye’ | 5 | ‘Eskirimo’ | 6 |
|  | ‘Keeyi’ | 6 | ‘Fatate’ | 3 | ‘Ganabasi’ | 1 |
|  |  |  |  |  | ‘Bondi’ | 8 |
|  |  |  |  |  | ‘Israeli’ | 14 |
|  | ***n=3*** | S =17 | ***n=3*** | S =20 | ***n=5*** | S=32 |
| Shannon |  | H’=1.09 |  | H’=1.32 |  | H’=1.37 |

Key: H’=Diversity index, n= Number of varieties landraces, S = number of collected landraces

#### 4.3.4. Sorenson similarity index of the wheat landraces

The Sorenson similarity index was used to detect similarities of wheat landraces among the three sampled kebeles of the study area. More similarity coefficient (0.33) was observed in the group of landraces at Digalubora and Burkitu kebeles. Less similarity index (0.25) was observed between Gushatemala and Burkitu (Digalubora) kebeles (Table 5).

**Table 5.** Sorenson similarity index of the wheat landraces of the study area

| Kebeles | Gushatemala | Digalubora | Burkitu |
| --- | --- | --- | --- |
| Gushatemala | 1.00 |  |  |
| Digalubora | 0.25 | 1.00 |  |
| Burkitu | 0.25 | 0.33 | 1.00 |

#### 4.3.5. Correlation analysis between wheat landraces distribution and farmland sizes

The distribution of wheat landraces and size of farmlands of the study area was 78% related and the relationship is a strong positive and significant (p=0.01) (Table 6).

**Table 6.** Correlation analysis of wheat landraces distribution and farmland sizes

| No_ | Distribution of wheat landraces | size of farmlands in hectare | r | p |
| --- | --- | --- | --- | --- |
| 1 | 8 | 0.25 | **0.78** | **0.01** |
| 2 | 10 | 0.5 |  |  |
| 3 | 16 | 1 |  |  |
| 4 | 13 | 1.5 |  |  |
| 5 | 22 | 2 |  |  |

Regarding the correlation analysis of some agronomic features for wheat landraces, the stem length and maturity time had a significant positive correlation (p≤0.01), the productivity in quintal per hectare was also positively correlated with maturity time and significant at (p≤0.05). However, productivity per hectare was negatively corrected with stem length (Table 7).

**Table 7.** Correlation analysis of agronomic features for wheat landraces

| variables | MT | SL | PrH |
| --- | --- | --- | --- |
| Maturity Time (MT) | 1.00 |  |  |
| Stem Length (SL) | 0.17 | 1.00 |  |
| Productivity/ kuntal/ Hectare (PrH) | 0.40 | -0.11 | 1.00 |

#### 4.3.6. Vernacular name and agronomic characteristics of wheat landraces

Farmers described wheat landraces based on phenotypic traits (example, seed color, and plant height), maturity time, end-use quality, and agronomic characteristics. For instance, white-colored wheat seed (example, “Israeli”); purple/red colored varieties seed (example, “Keeyi”); black colored varieties of wheat seeds (example, ‘Qamadiguracha’) but ‘Fatate’ have the mixture of white and black colored wheat seeds. Furthermore, wheat landraces reached maturity between 4-6 months, and they were purposively used for different end-use purposes (examples, for making bread, ‘injera’, gruel, and beverage, etc.) (Table 8).

**Table 8.**
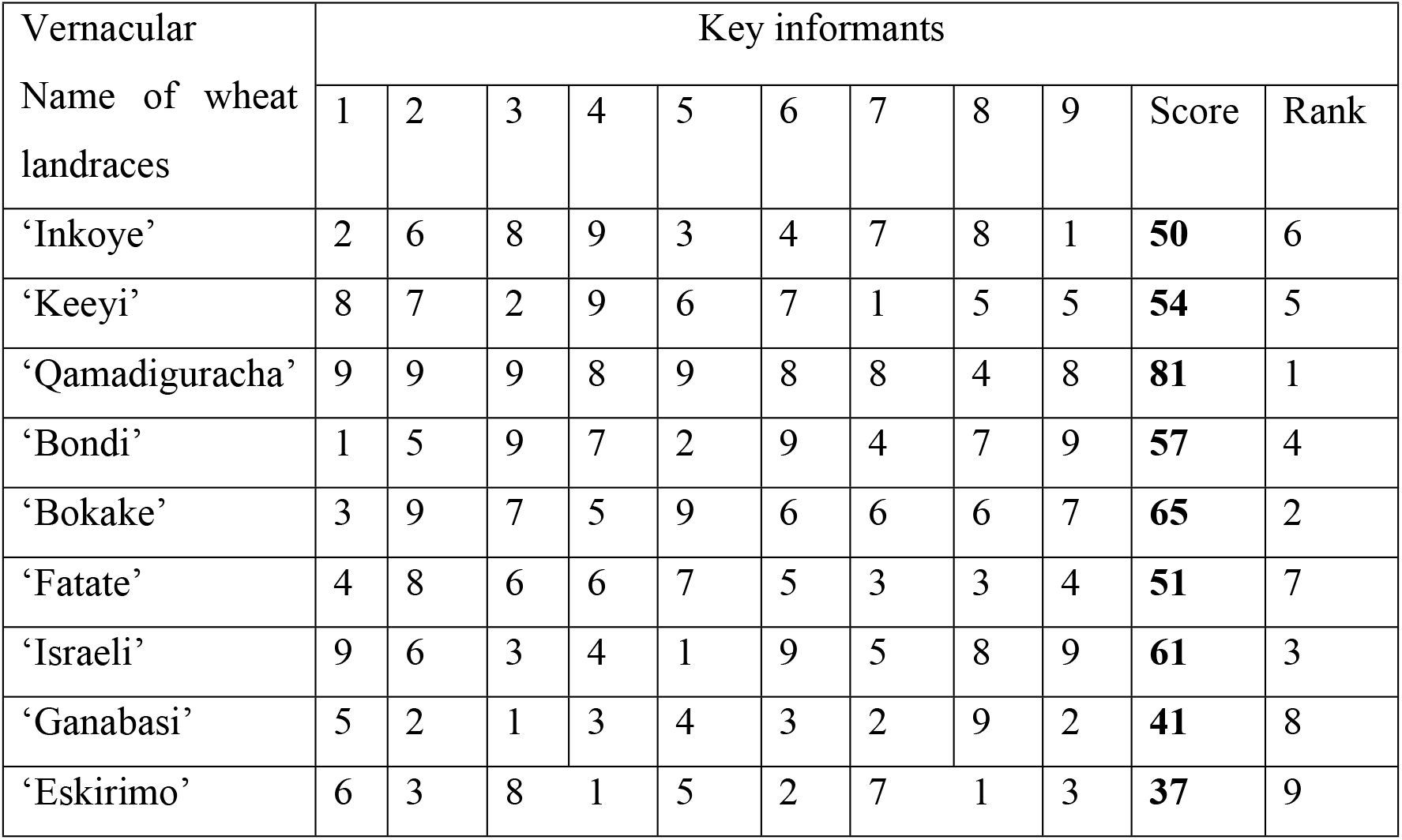
Preference ranking for wheat landraces in their end-uses (1 −9): (9, for most Preferred and 1, for least preferred)

‘Qamadi guracha’ has black seeds and long hulls in phenotypic (Fig.4). Its introduction year was not known and reached maturity at 5 months into the study area. According to the farmers, these landraces are the most drought, pests, birds, and disease resistant. Additionally, farmers ‘Qamadi guracha’ is most preferred for making bread, ‘injera’, gruel, and beverage. Furthermore, it has good qualities of straw and shelf life than other wheat landraces (Table 8).

**Fig.4.**
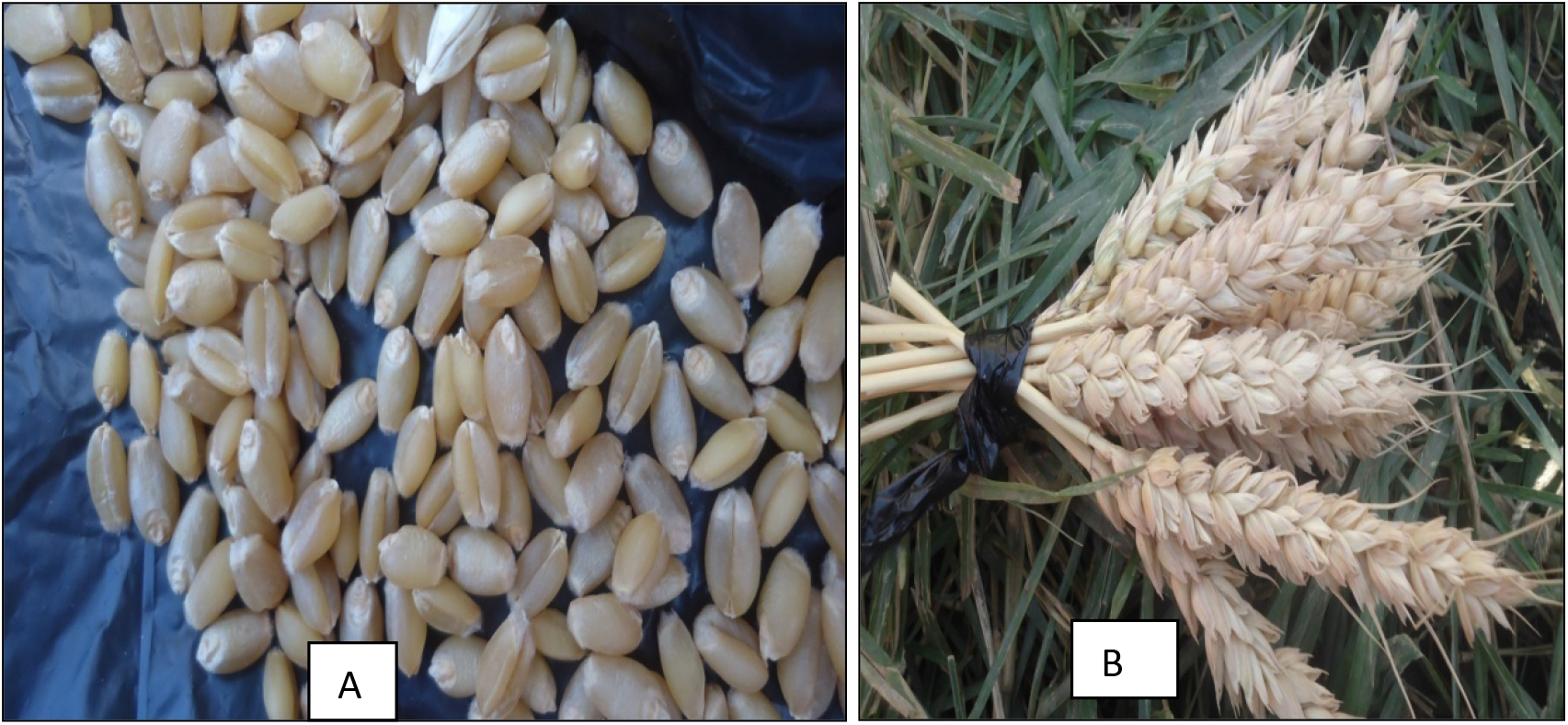
‘A’ seeds of Isira’el and ‘B’-Israeli with hulls

‘Bondi’ has hulls, white seeds, and no spikes (Fig.5). Its introduction year into the study area was not known. According to respondents, ‘Bondi’ can tolerate drought and low soil fertility conditions; therefore, it gives better yields than other landraces such as ‘Qamadi guracha’. The flour of ‘Bondi’ is used for making ‘injera’ and bread of good taste and quality, and homemade porridge (Appendix 1).

**Fig.5.**
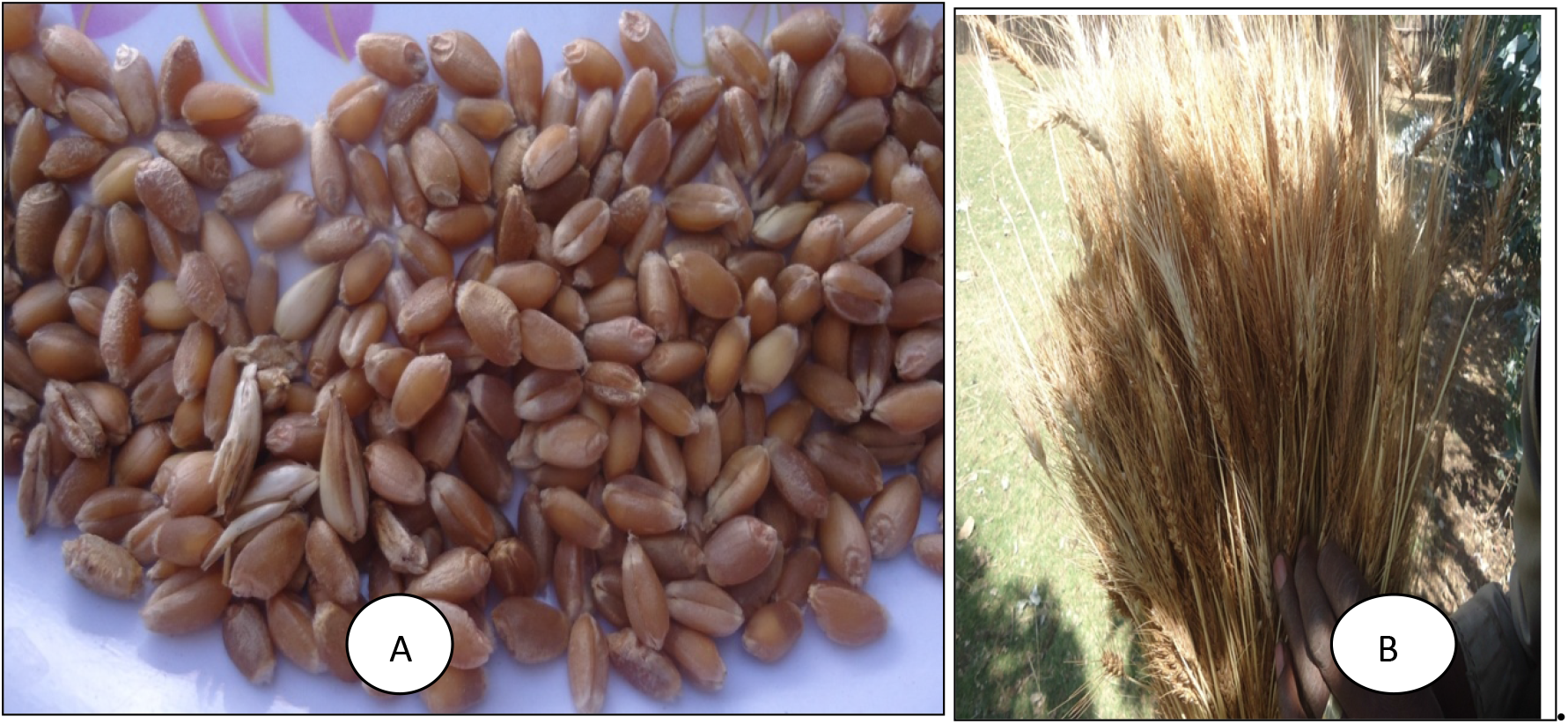
’A’-seeds of ‘Keeyi’ and ‘B’- ‘Keeyi’ with hulls.

‘**Israeli**’ is non-spikes, white wheat has big seeds and a similar plant stands as ‘Bondi’. As a result, some farmers had confused this landrace with ‘Bondi’ and only experienced farmers could discern the difference between the two landraces (Fig. 6). It was introduced into the study area in 1967. According to respondents, agronomically, this landrace requires fertile soils; susceptible to drought and cold and it loses easily the hulls as drying. It reached maturity in five months. Additionally, bread made from ‘Israeli’ is regarded by the farmers as good as that made from modern wheat. Its grains were also preferred for making ‘aka’i or kollo’ in the study area (Appendix 1).

**Fig.6.**
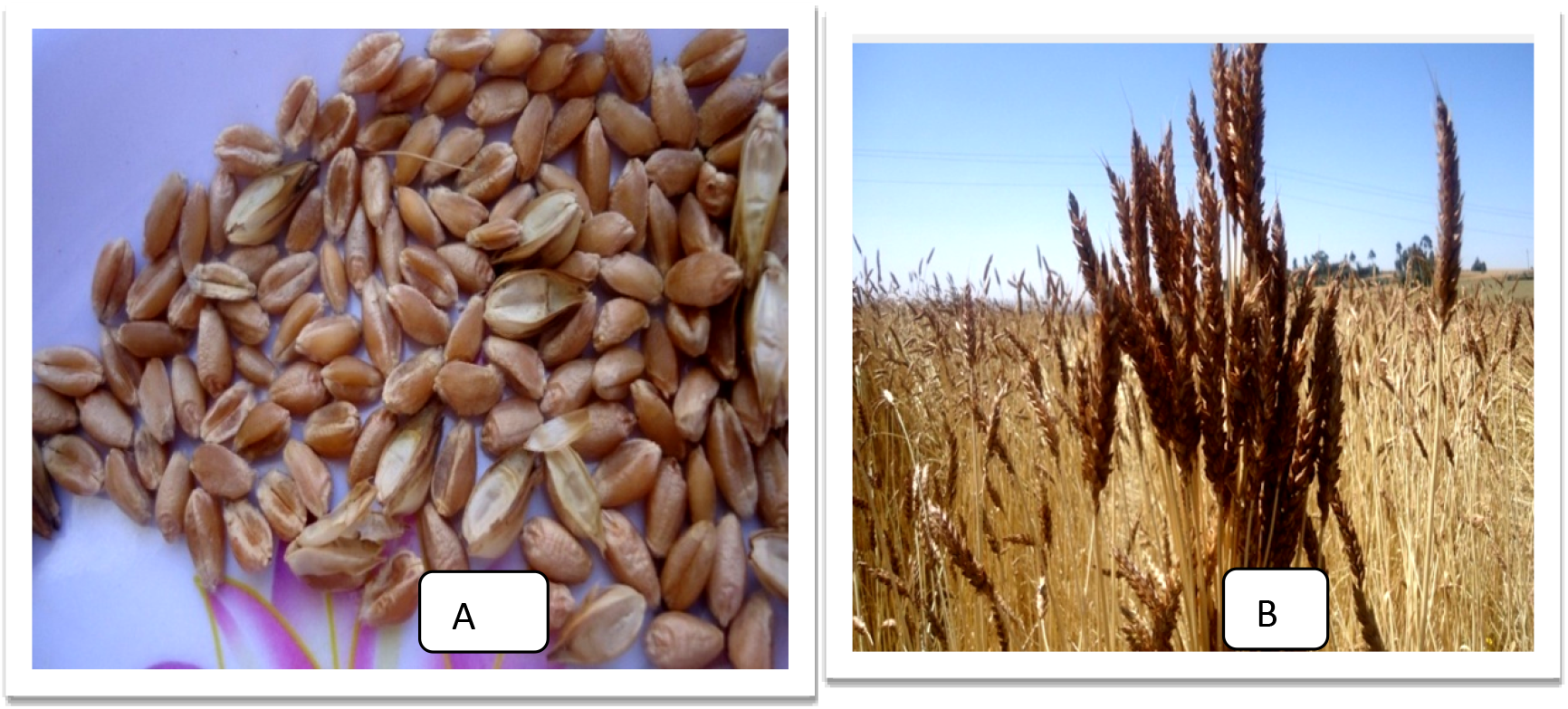
‘A’ Eskirimo red seed; ‘B’ straw of Eskirimo

**‘Keeyi’,** which is also known as ‘Qamadi dima’ meaning ‘red –wheat’, has hulls and small spikes (Fig.7). According to the farmers, this landrace was introduced into the study area in 1983. Concerning the agronomic characteristics of ‘Keeyi’, it was widely adapted to disease, drought, pests, and low soil moisture; it is usually grown at the end of the rainy season on residual moisture. Moreover, ‘Keeyi’ was used for making the beverage, ‘injera’ and bread in the stud area (Table 8).

**Fig.7.**
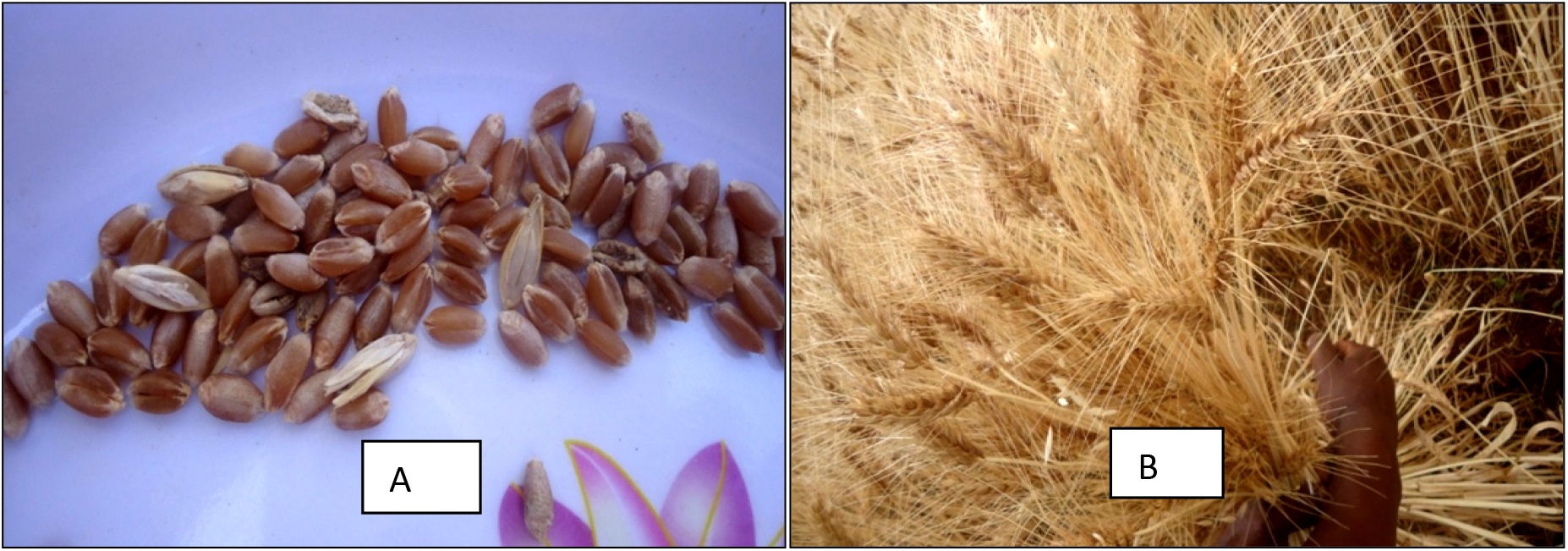
‘A’ Inkoye red seeds; ‘B; Inkoye seed with hulls wheat landrace was also highly preferred for making bread, ‘injera’, and beverage (Table8).

**‘Eskirimo’** is non-spikes, has red seed and long stem length which measured about 59.9cm (Fig.8). It was introduced into the study area in 1979. Farmers also responded that this wheat landrace was tolerant to drought and lodging; it had a long maturity period of about six months and yielded about 24.36 quintals per hectares per year. ‘Injera’ made from ‘Eskirimo’ was regarded by the farmers as good as that made from barely. Its flour was also preferred for making bread in the study area (Table 15).

**Fig.8.**
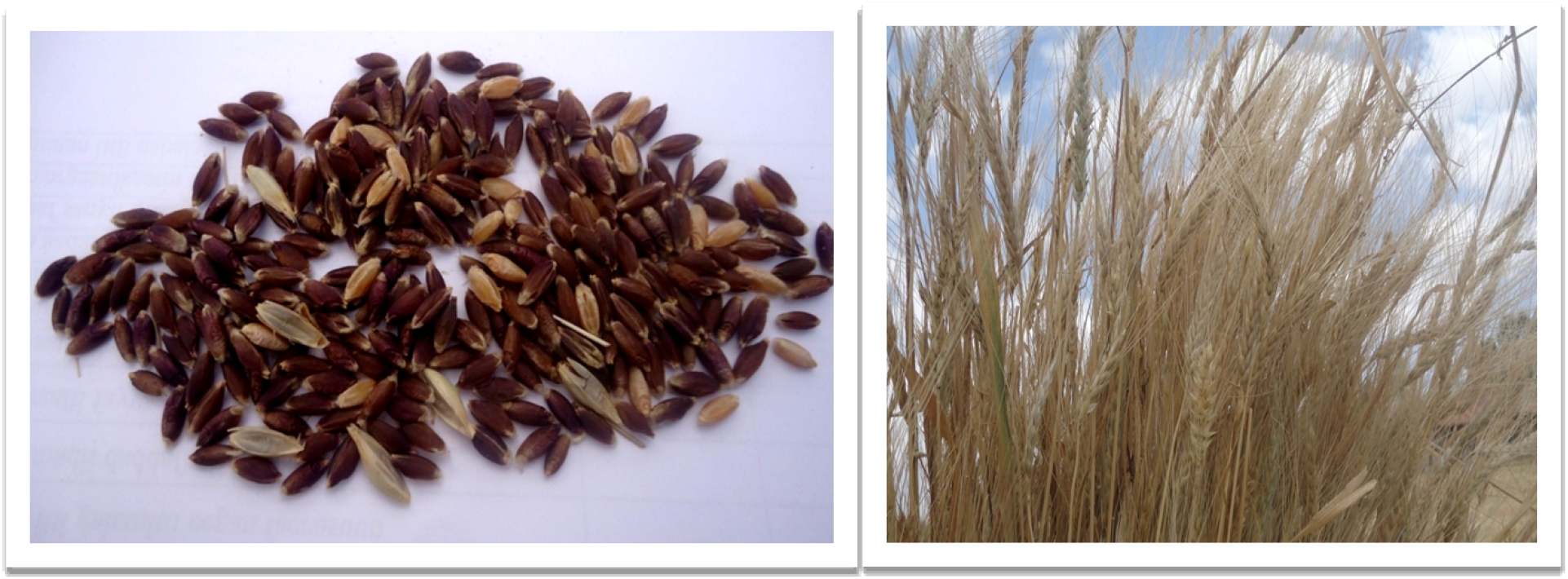
‘A’- ‘Fatate’ seeds; ‘B’-Fatate with straw

‘**Inkoye’,** which is also red–seeded, has hulls (Fig.9). It was introduced into the study area in 1959 E.C. The Farmers, DA workers, and district Agricultural Experts reported that ‘Inkoye’ had better adaptation to lodging, low fertile soils, diseases, and cooler temperatures in the study area. Its maturity time, average stem length, and productivity per hectare per year were five months, 64cm, and 32kuntals/hectare respectively. It was mainly used for the preparation of bread, beverage, ‘injera’, and rarely for making gruel (Table 8, Table 15).

**Fig.9.**
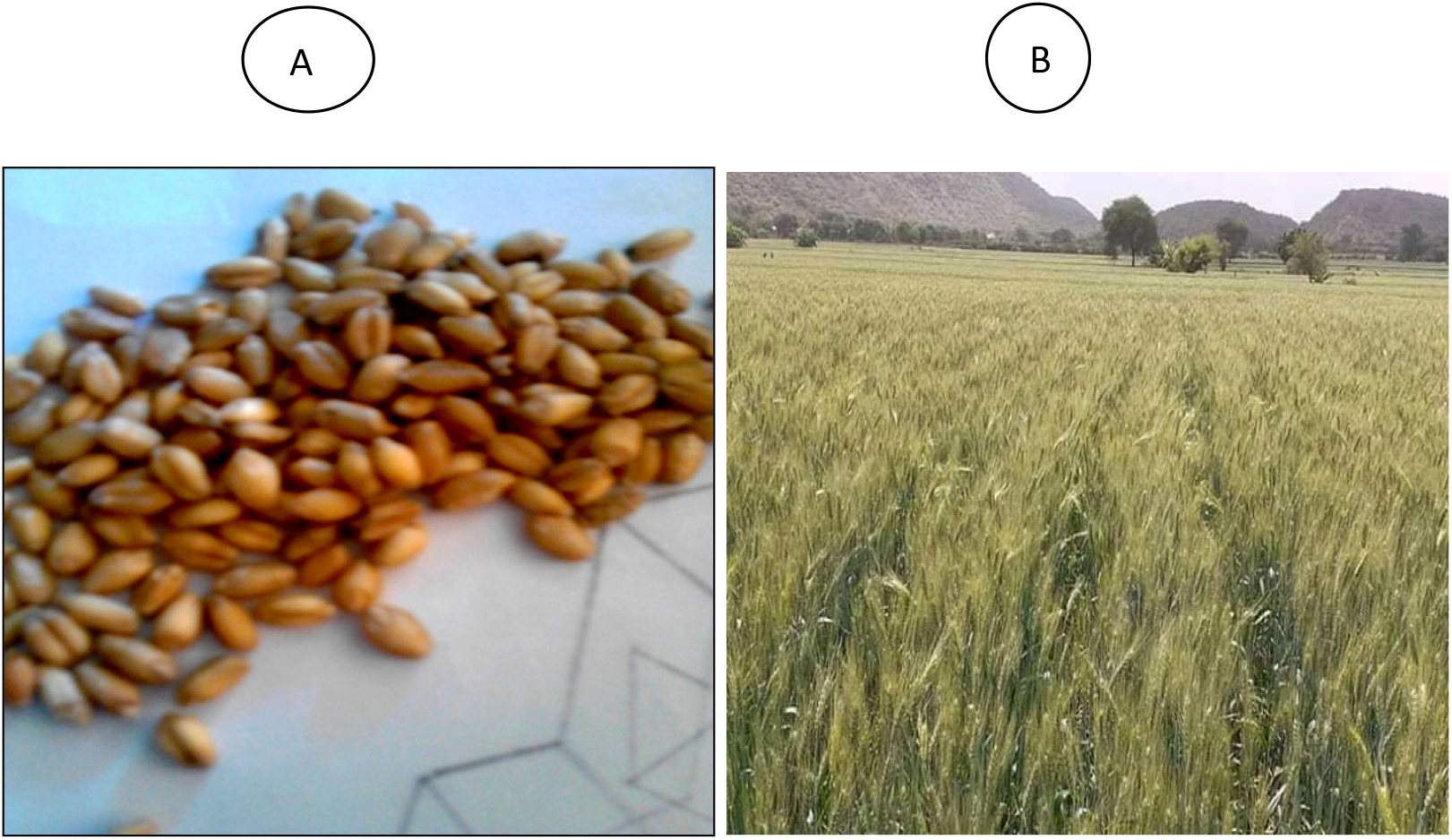
Seed of ‘Ganabasi’ wheat landrace

**‘Fatate’ (‘Sergagna’),** which is the black and white seeded, has hulls and long spikes (Fig.10). Most farmers of the survey area reported that ‘Fatate’ can reach maturity in five months and is mainly grown at high altitude areas of the study area. It could resist lodging and pests and need black soil. The straw of ‘Fatate’ was used for traditional roofing and fodder for animals. This wheat landrace was also highly preferred for making bread, ‘injera’, and beverage (Table 8).

**‘Ganabasi’,** which is the wheat landrace, has white seeds (Fig.10). Most farmers of the survey area reported that ‘Ganabasi’ could reach maturity in five months. Nevertheless, farmers from Burkitu reported that this landrace needs only about four months until maturity presumably because of lower altitude, warmer temperature, and faster development in the kebele. The fertile soil was required for better yields. Despite its height, lodging is not a problem. ‘Ganabasi’ was highly preferred for making ‘injera’ and beverage in the study area (Table 8).

**‘Bokake’**, which is blackish-seeded, has short spikes and red-hulls. It was introduced into the study area in 1964. The majority of the farmers of the study area reported that ‘Bokake’ reached maturity in five months. It could resist lodging and pests. It needs black soil and be sowed with barely. It was used for making bread, ‘Kollo’, beverage, and gruel (Table 8).

#### 4.3.7. Preference ranking of wheat landraces in terms of end-use purposes

Key informants ranked the nine wheat landraces based on their end-use purposes by giving 9 for most valuable and 1 for least valuable. The scores given to each landrace as per informant preference were added and ranked. Consequently, ‘Kamadiguracha’ was ranked first and ‘Eskirimo’ preferred least in terms of end-use purposes (Table 9)

**Table 9.**
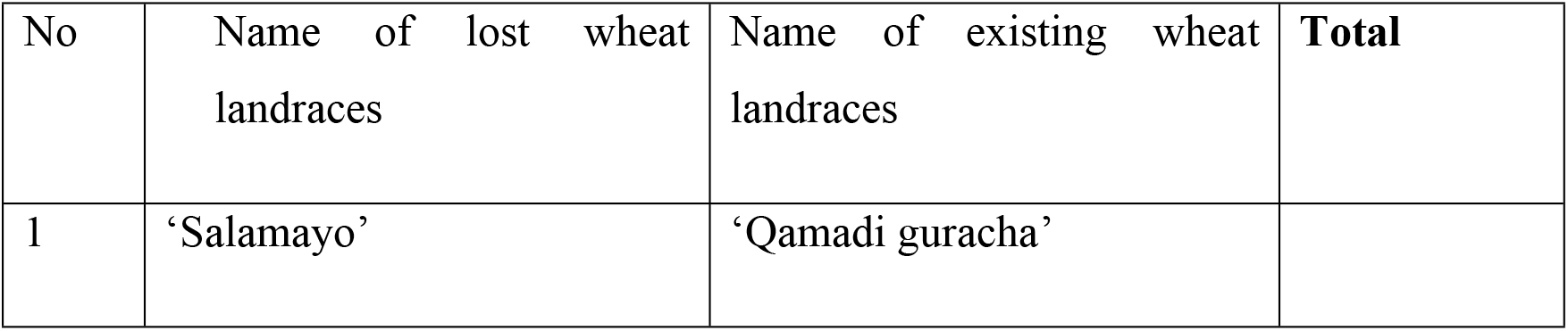
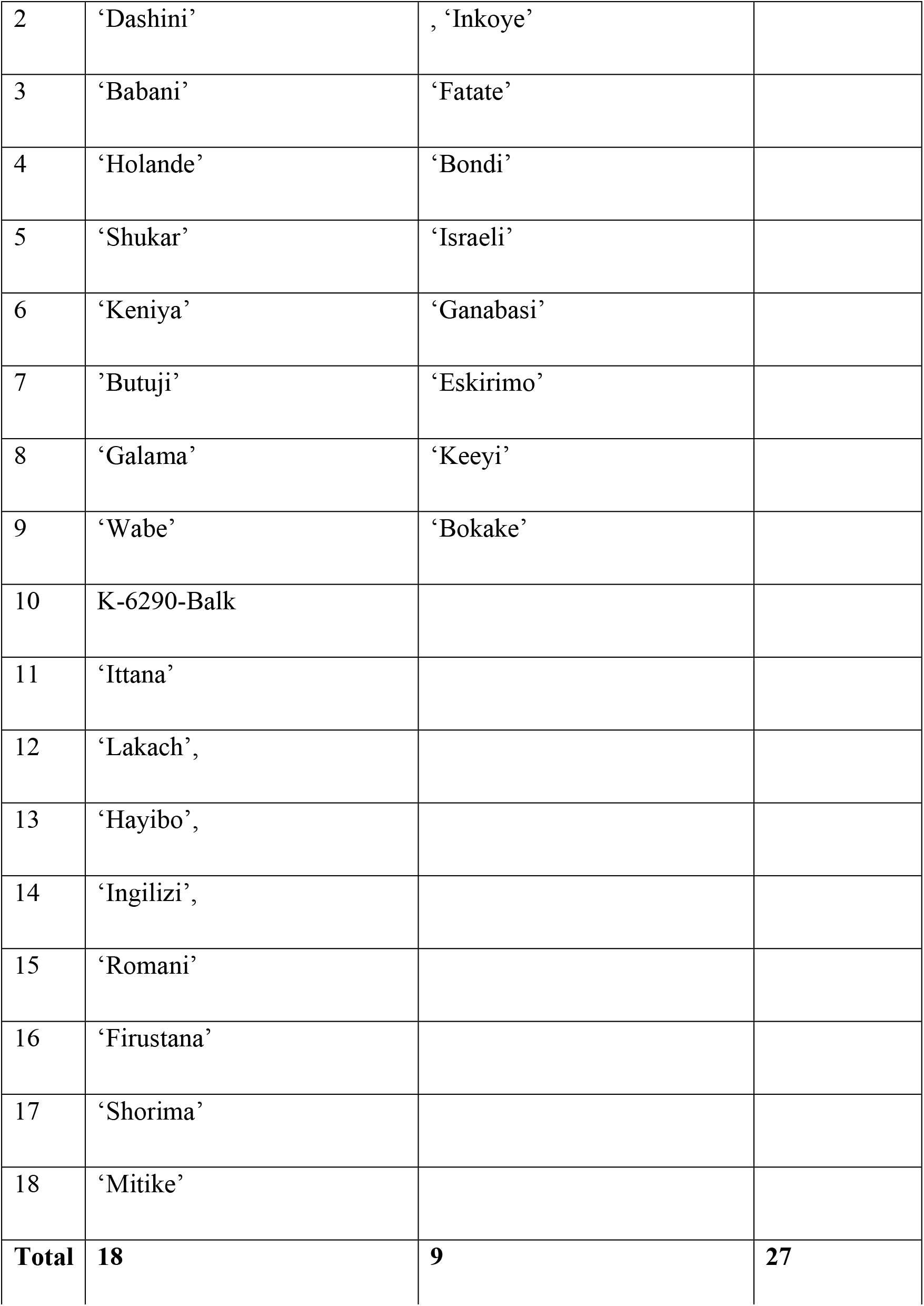
List of lost and existing wheat landraces

| No | Name of lost wheat<br>landraces | Name of existing wheat<br>landraces | Total |
| --- | --- | --- | --- |
| 1 | ‘Salamayo’ | ‘Qamadi guracha’ |  |
| 2 | ‘Dashini’ | , ‘Inkoye’ |  |
| 3 | ‘Babani’ | ‘Fatate’ |  |
| 4 | ‘Holande’ | ‘Bondi’ |  |
| 5 | ‘Shukar’ | ‘Israeli’ |  |
| 6 | ‘Keniya’ | ‘Ganabasi’ |  |
| 7 | ’Butuji’ | ‘Eskirimo’ |  |
| 8 | ‘Galama’ | ‘Keeyi’ |  |
| 9 | ‘Wabe’ | ‘Bokake’ |  |
| 10 | K-6290-Balk |  |  |
| 11 | ‘Ittana’ |  |  |
| 12 | ‘Lakach’, |  |  |
| 13 | ‘Hayibo’, |  |  |
| 14 | ‘Ingilizi’, |  |  |
| 15 | ‘Romani’ |  |  |
| 16 | ‘Firustana’ |  |  |
| 17 | ‘Shorima’ |  |  |
| 18 | ‘Mitike’ |  |  |
| **Total** | **18** | **9** | **27** |

### 4.4. On-farm genetic erosion and its causes of wheat landraces

#### 4.4.1. Extent of on-farm genetic erosion of wheat landraces

Many farmers (97.8%) and DA workers replied that the number of landraces has been decreasing starting from the last 20 years up to 2017. As a result, from a total of 27 wheat landraces that were listed by farmers, 18 of them were lost and they were reduced to 9 landraces which are being cultivated in the site presently (Table 9).

#### 4.4.2. Factors responsible for the genetic erosion of wheat landraces

In the present study, about 67(50%), 34(25.4%), 17(12.2%), and 10(7.5%) of farmers responded the majority of wheat landrace had been lost due to four main factors: relative low productivity of wheat landraces, the introduction of modern wheat variety or hexaploid wheat (example, ‘Hidase’ and ‘Digalu’), the introduction of other more productive crops (example, barley, and maize), and land degradation/ soil infertility/ respectively (Table 10). In addition to this, the document analysis also showed that more frequent occurrences of cold temperatures and changes in market prices were the factors that might cause the loss of wheat landraces from the survey site.

**Table 10.** Genetic erosion of wheat landraces in the study area

| Summary of genetic erosion of wheat landraces |  |
| --- | --- |
| Number of landraces recalled by the farmers grown before 2 decades | 27 |
| Number of landraces grown during 2016 to 2017 | 9 |
| Genetic integrity (%) | 33.3 |
| Genetic erosion (%) | **66.6** |

**Table 11.** Factors that cause the genetic erosion of wheat landraces

| Item | Causes | Respondents | percentage |
| --- | --- | --- | --- |
| What are the causes for reduction of what landraces ? | Land degradation/soil infertility | 10 | 7.5% |
|  | Introduction of Modern wheat | 34 | 25.4% |
|  | Introduction of other more productive crops | 17 | 12.2% |
|  | Diseases | 1 | 0.75% |
|  | Climate change | 4 | 2.98% |
|  | Low productivity of landraces | 67 | 50% |
|  | Drought | 2 | 1.50% |
|  | **Total** | **134** | **100%** |

### 4.5. Factors that contribute to on-farm conservation of wheat landraces

About 46%, 28%, and 26 % of farmers responded that the food quality, market value, and resistance to a pest, birds, and diseases of wheat landraces were the major factors that made them preserve the wheat landraces on their farmlands respectively (Table12). Additionally, document analysis indicated that the big seed size, long shelf life, and straw quality of some landraces were moderate incentives that made some farmers conserve them.

**Table 12.**
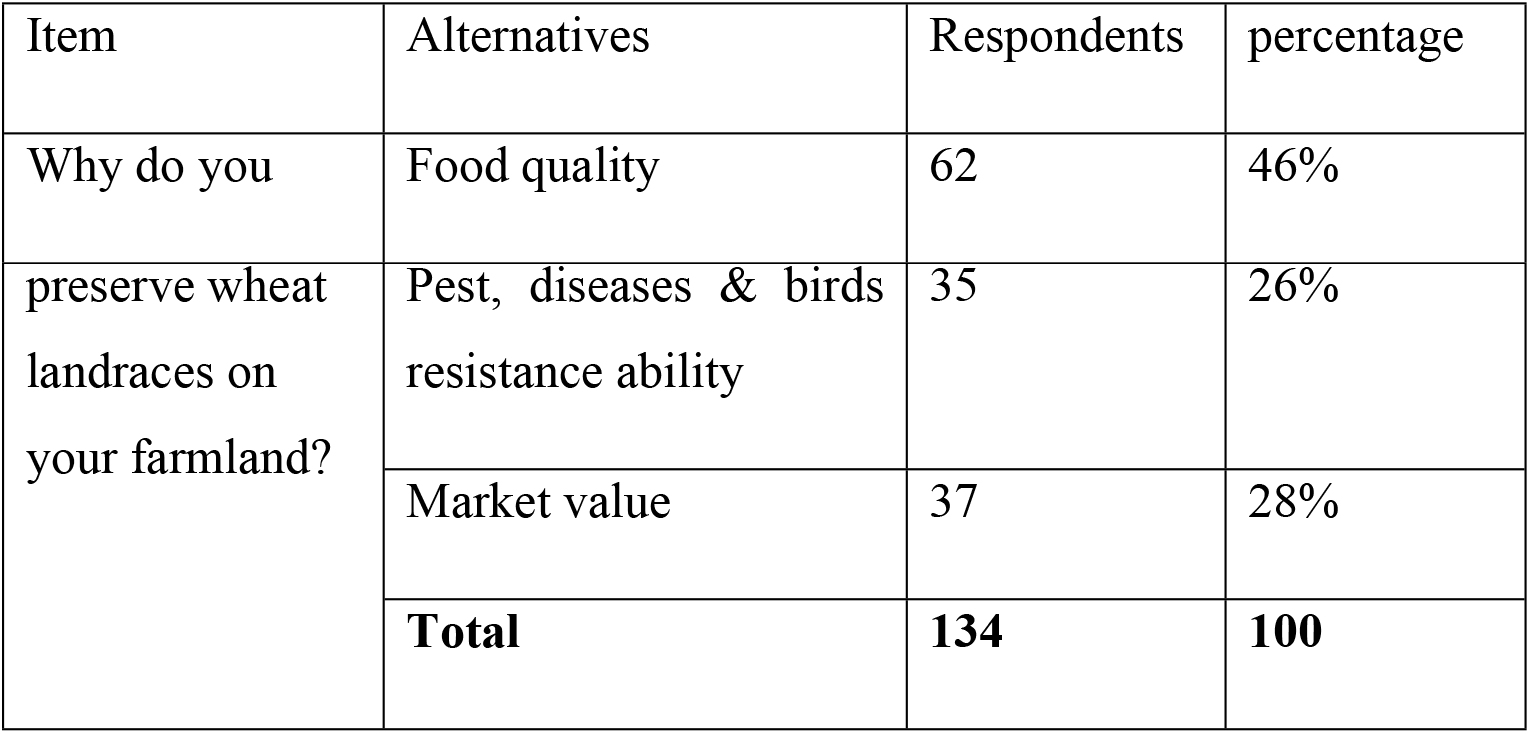
Factors to preserve the wheat landraces on farmlands by farmers

| Item | Alternatives | Respondents | percentage |
| --- | --- | --- | --- |
| Why do you | Food quality | 62 | 46% |
| preserve wheat landraces on your farmland? | Pest, diseases & birds resistance ability | 35 | 26% |
|  | Market value | 37 | 28% |
|  | **Total** | **134** | **100** |

### 4.6. Major bottlenecks influencing the conservation of wheat landraces

According to the 52(38.8%), 31(23.1%), 23(17.2%), and 21(15.6%) of respondents, seed selection system, insufficient crop yields, insufficient land holding, and soil infertility were major bottlenecks that hindered the conservation of wheat landraces in the study area respectively(Table13).

**Table 13.** Bottlenecks influencing the conservation of wheat landraces

| Item | Alternatives | Respondents | percentage |
| --- | --- | --- | --- |
| major bottlenecks | Insufficient land holding | 21 | 15.6% |
|  | Seed selection system | 52 | 38.8% |
|  | soil infertility | 23 | 17.2% |
|  | insufficient crop yield | 31 | 23.1% |
|  | farmland management system | 3 | 2.2% |
|  | rainy season problems | 4 | 2.9% |
|  | **Total** | **134** | **100** |

### 4.7. Farmers’ roles to conserve wheat landraces for future

According to the respondents, the major and least factors contributing to the future conservation of wheat landraces were re-sowing on farmlands using organic fertilizers (manure, **compost, etc.) and on-farm seed selection respectively (Table 14).**

**Table 14.** Roles of farmers to conserve wheat landraces in the study area

| Roles of farmers | Respondents | percentage |
| --- | --- | --- |
| On-farm seed selection | 2 | 1.5% |
| On-farm conservation by re-sowing | 57 | 42.5% |
| Protecting from pests traditionally | 11 | 8.2% |
| Harvesting on maturity times | 15 | 11.2% |
| Using synthetic fertilizers | 3 | 2.23% |
| Using organic fertilizers | 5 | 3.75% |
| Saving/managing seeds for future | 38 | 28.4% |
| **Total** | **134** | **100%** |

**Table 15.**
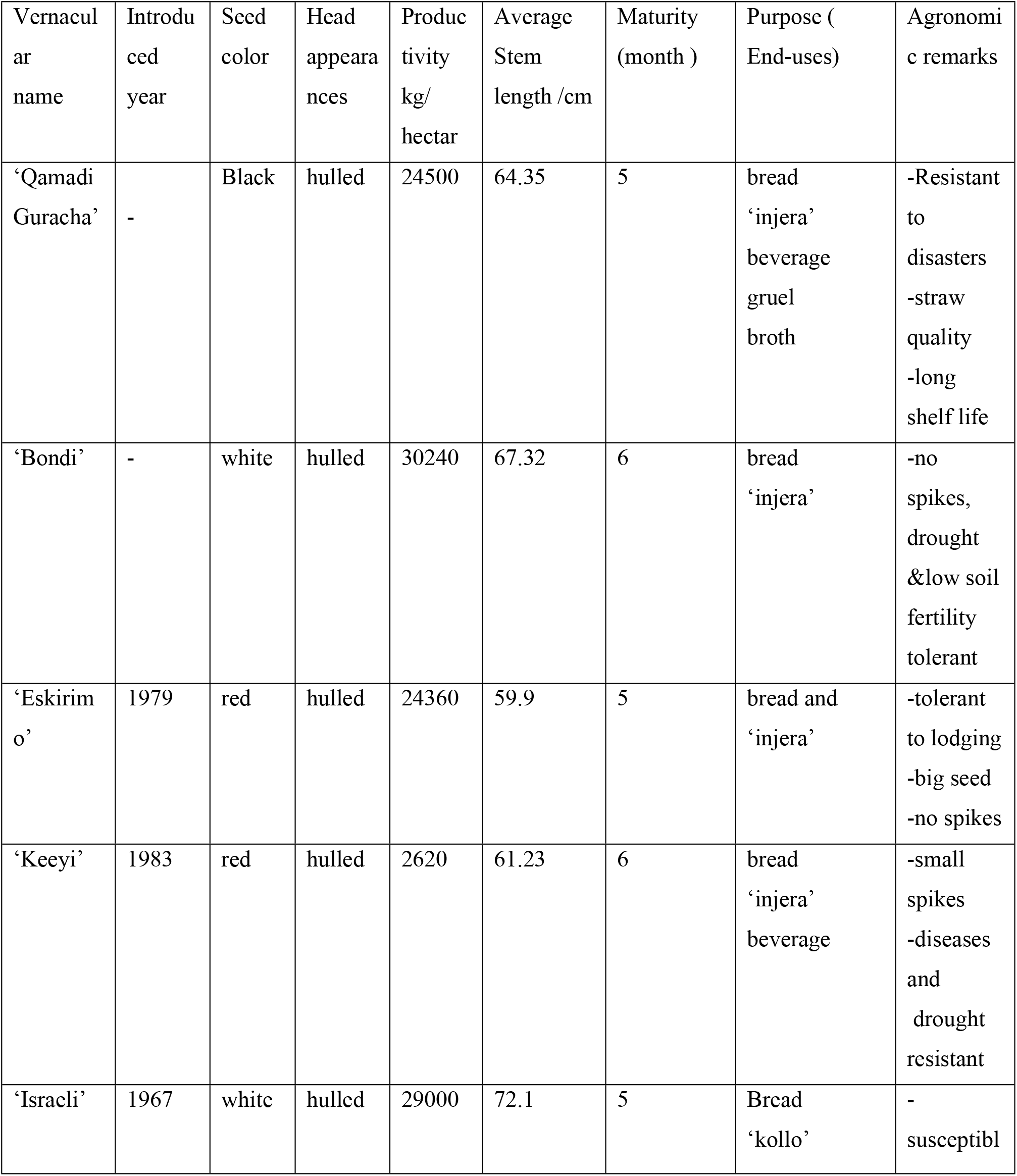
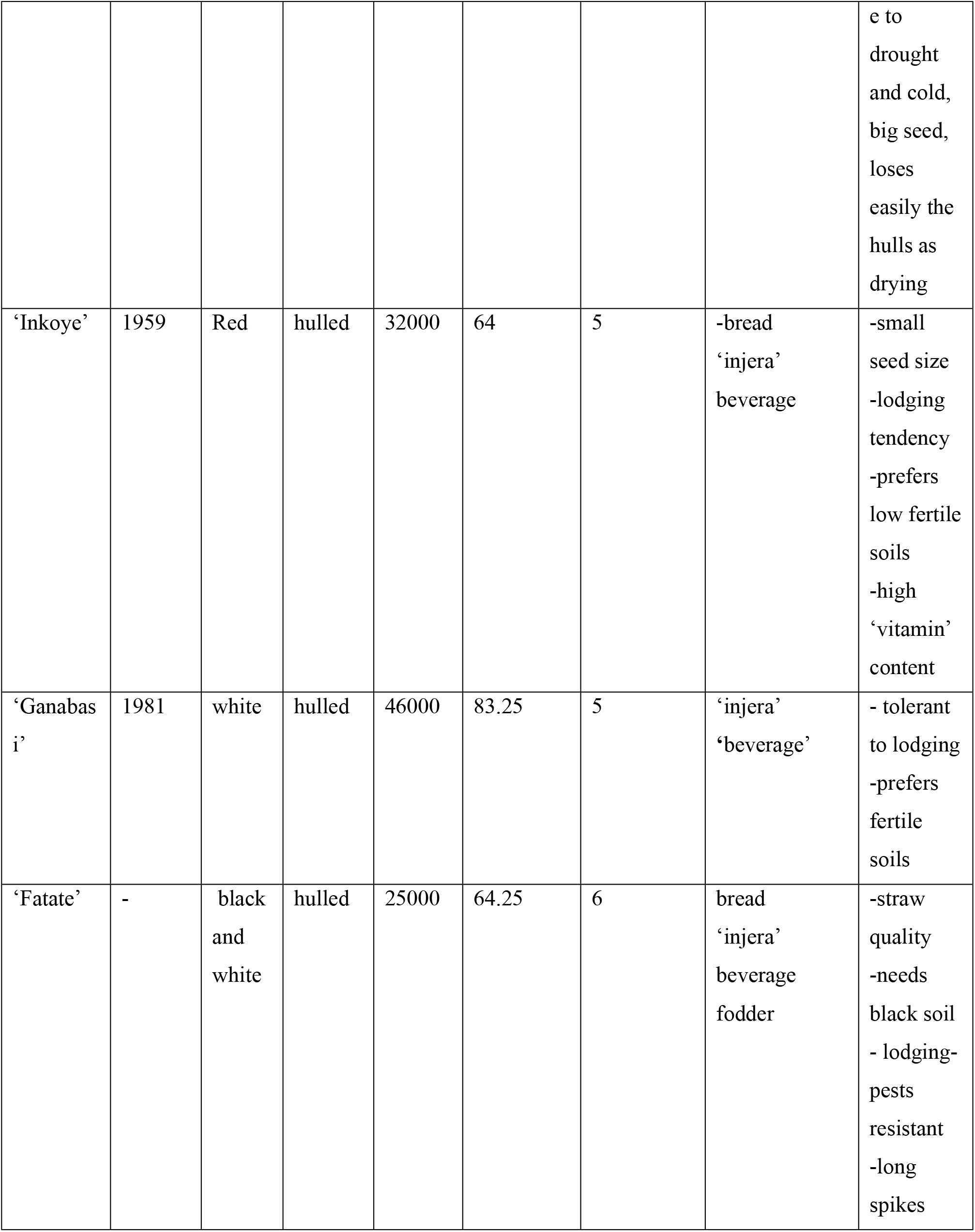
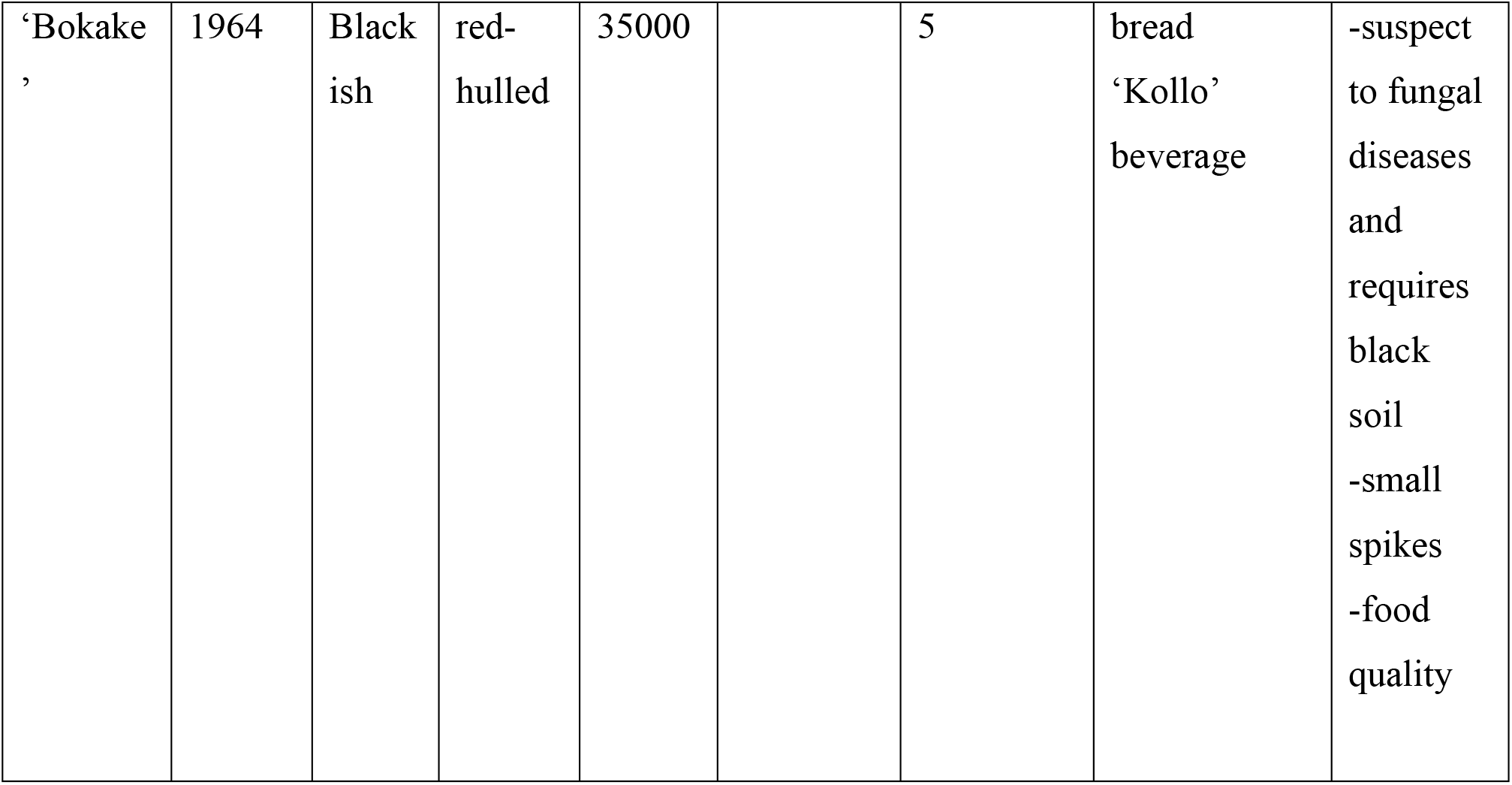
Vernacular names and agronomic features of wheat landraces identified in this study

### 4.8. Naming of wheat landraces

Farmers in the study area used to name the wheat landraces varieties by using seed color, maturity time, and food quality which have their meanings. For instance, ‘Kamadi guracha’ (Afaan Oromo) was named after its black seed color. ‘Inkoye’ was named after its very small seed, usually has good food quality. ‘Ganabasi’ (Afaan Oromo) was named as ‘winter - crosser’ meaning it could be used in the winter season due to its surplus. ‘Bokake’ (Afaan Oromo) was named after it ‘expands’ while its seeds are cooked. Likewise, ‘Fatate’ (Amharic) means a mixture of white and black seeds of wheat landraces. Bondi (Afaan Oromo) was named after its market value which could buy ‘boundi or birr’.

## 5. Discussion

In a total, 27 wheat landraces were reported to be grown two decades up to now. Of these, only 9 of wheat landraces are still cultivated by farmers. Among these available landraces, ‘Qamadi guracha’ is the most common in the study area. The computed Shannon diversity index of currently growing wheat landraces ranges from 1.09 to 1.37 among the groups. This shows that there was a highly diverse of wheat landraces at present in the study area relatively. The estimated genetic erosion for wheat landraces was found to be 66.7% due to major factors: introduction of modern wheat varieties; the introduction of other more productive crops; landraces low productivity; and infertility of soil. Food quality, Pests (for example, birds and insects), diseases resistance, market value, and straw quality were factors that initiated the farmers to maintain the genetic diversity of landraces. However, the Preservation of these landraces is influenced by bottlenecks like seed selection systems, soil infertility, and insufficient crop yields.

### 5.1. On-farm genetic diversity of wheat landraces

In the present study, in total 27 wheat landraces were identified that had been cultivated in the past 20 years to the present from the sampled kebeles in Digalu and Tijo district of Arsi Zone. Similarly, Faris Hailu (2011) indicated that there was a rich diversity of wheat landraces, which contained 15 to 37 varieties, in the Arsi highlands. About 26 wheat landraces were nearly identified by Tesfaye Tsegaye and Berg (2007) from Akaki and Lume Districts of East Shoa, and Negash Geleta and Grausgruber (2013) also discovered 20 different landraces from Ambo and Dandi of West Shoa. Furthermore, 21 durum wheat landraces, 11 of which were collected from North Shoa and the remaining 10 from Bale (Fassil Kebebew *et al*., 2001).

The Shannon diversity index of wheat landraces was estimated based on the number of varieties collected. Accordingly, the Gushatemala kebele showed the highest diversity (H’= 1.37) followed by Digalubora kebele (H’= 1.32). Burkitu kebele was found to be less diverse in terms of the number of varieties collected in this study area (H’= 1.09). Similarly, Brown (1983) indicated that the diversity estimated based on several wheat landraces (Shannon diversity index), Arsi has the higher diversity index (H’=1.31) than other Ethiopian wheat-producing areas such as North Shoa.

Sorenson similarity index among wheat landraces of the study area ranged from 0.25 - 0.33 which are not above 0.5. This indicated that there were low similarities in wheat landraces composition among the study kebeles; and also implied that there werewash wheat landraces diversity in Digalu and Tijo district of Arsi Zone. The existence of low similarities among wheat landraces implied that all the kebeles are important in terms of the diversity of wheat landraces. This finding can be confirmed that the low similarity index existence among landraces indicated that ddiversityof wheat landraces and related to dissimilar habits for growing and human selection on landraces (Kent and Coker, 1992). On the other hand, the correlation analysis of the distribution of wheat landraces and the size of farmlands of the study area was 78% correlated. The relationship was acceptable and positive because the ‘r-value is between 0.7 and 0.8. According to Zawdie Bishaw (2014), a positive association is expected between large-sized farmlands and the number of wheat landraces because on wider farmlands farmers can produce more wheat landraces.

The correlation analysis of some agronomic features (between stem length and maturity, productivity in kg per hectare and maturity time, productivity in quintal per hectare, and stem length) for wheat landraces was done. In this line, the stem length and maturity time had a significant positive correlation (p≤0.01). In the same way, productivity in kg per hectare was positively correlated with maturity time and significant at (p≤0.05). This finding can be confirmed that the longer maturity time, the higher grain yield of wheat landraces which indicates the positive correlation between the two (Zawdie Bishaw *et al*. 2014). However, productivity in quintal per hectare was negatively corrected with stem length and number of wheat landraces. A similar finding was reported by Getachew Belay *et al*. (1993) that the grain yield is negatively correlated with the average stem length of Ethiopian wheat landraces.

### 5.2. Agronomic characteristics of wheat landraces

The study showed wheat landraces with diverse morphological features (e.g. seed color, stem length, and head appearance). For example, landraces such as ‘Inkoye’, ‘Eskirimo’, and ‘Keeyi’ have red/purple seed color whereas ‘Kamadiguracha’ and ‘Bokake’ have Black/blackish colored seeds but ‘Fatate’ is the mixture of white and black colored seeds. Similarly, Zeven (2002), in the study conducted in the central and southeastern highlands of Ethiopia, indicated that a purple color seed was originally only found in cultivated wheat landraces of Ethiopia. Some wheat landraces have no spikes whereas others have very long spikes; some are big-seeded but others have small seeds. Moreover, there are specific endemic characteristics for some of the tetraploid wheat species in Ethiopia, such as big grain, spikeless or with spikes, and beardless or half-bearded hard durum wheat (Jain, 2000; Teklu Tesfaye and Hammer, 2006).

On the other hand, the wheat landraces exhibited differences in other agronomic characteristics. Some were adapted to less fertile soils, whereas others need fertile soil; others exhibited differential responses to diseases, pests (for example, insects and birds), adaptation to soil infertility, and lodging. This is similar to the idea stated as “in history, Ethiopian tetraploid wheat landraces, disease, pest, drought resistance and other stresses, adaptation to low soil fertility and other characteristics are the useful agronomic remarks exhibited by them” (Melaku Worede, 1997). Tesfaye Te mma *et al*. (1993) also stated that Ethiopian wheat landraces are usually mixtures of different agro-types having wider gene pools within one population that make them to adapt the change of climatic and edaphic factors.

Moreover, Key informants ranked the nine wheat landraces based on their end-use purposes by giving 9 for most valuable and 1 for least valuable in the study area. In this line, ‘Qamadi guracha’ was ranked first and ‘Eskirimo’ preferred least. This indicated that ‘Qamadi guracha’ is the most valuable in making end-use products such as bread, ‘injera’, beverage, gruel, and broth in the study area. This finding re-affirmed that some landraces such as ‘Tikur sende’ are used for a specific purpose such as for brewing local beer or spirits; as a result, therefore, it can be ranked at the top of others in North Shoa of Amhara region (Efrem Bechere *et al*., 2000; Zawdie Bishaw, 2014).

### 5.3. The Extent of Genetic erosion of wheat landraces

In the current study, the number of wheat landraces has been decreasing from the last 20 years up to this year (2017) from 27 to 9 in the study area. Consequently, the overall on-farm loss (genetic erosion) of wheat landraces diversity in the survey area was found to be 66.6% (Table 10). A similar scenario was indicated by Negash Geleta and Grausgruber (2013) from West Shoa that the genetic erosion of landraces found to be 75.0% and 61.5%, for Ambo and Dandi districts respectively; and Faris Hailu (2011) also discovered the overall genetic erosion in the in Ethiopia, 32.0%, 35.3%, 55.9%, 84.4%, and 84.0% erosion was found for *T. durum, T. turgidum, T. aethiopicum, T. polonicum*, and *T. dicoccon* respectively. In this study, the estimated level of genetic erosion was 66.6% which is higher than species that are said to be genetically eroded (i.e., GE=23-43%) (IBC, 2001). Furthermore, Brush (1999) argued that a loss of diversity of landraces implies a big threat for reduction of the pool of genetic material available for breeding to enhance productivity and ensure environmental stability.

### 5.4. Factors responsible for the genetic erosion of wheat landraces

In this study, the major factors responsible for genetic erosion of wheat landraces were low productivity of landraces, the introduction of improved wheat varieties (example, ‘Digalu’ and ‘Hidase’), the introduction of other more productive crops (example, barley), and infertility of soil/land degradation. The previous study (Faris Hailu, 2011) reported that the most important factors for loss of landraces were reduction in land size(cultivated area), displacement by released/modern varieties of hexaploid wheat and teff, reduced benefit from the landraces, and displacement by other crops and chat in Ethiopia. Other factors like climate change and the long maturity time of landraces were also causes of genetic erosion of wheat landraces in Ambo and Dandi Districts (Negash Geleta and Grausgruber, 2013). In addition, drought is also one factor of genetic erosion in tetraploid wheat landraces, especially in the eastern part of Ethiopia (Teklu Tesfaye and Hammer, 2006; Berg and Efrem Bechere, 2007)

### 5.5. On-farm factors that make the farmers preserve the wheat landraces

In the present study, the major factors responsible for the continued cultivation of wheat landraces by the farmers were: food quality, pest resistance, and market value. Additional minor factors such as seed size, long shelf life, and straw quality were moderate incentives that made the farmers conserve the wheat landraces. Similarly, Negash Geleta and Grausgruber (2013) explained that many farmers initiated to conserve some wheat landraces might be due to their unique end-use quality and wide adaptation to changing environments that are obtained from wheat landraces but not from improved varieties. For these reasons, the majority of the farmers were committed to conserving the genetic diversity of wheat landraces for the future in *in-situ* by re-sowing on farmlands, saving/managing seeds for the future, and using organic fertilizers. Similarly, Firdissa Eticha *et al*. (2010) argued that sustained on-farm conservation, saving for the future and sustainable utilization will ensure the continuous conservation of wheat landraces in Ethiopia.

### 5.6. Bottlenecks influencing the conservation of wheat landraces

Insufficient land holding, a seed selection system, and insufficient crop yield were major bottlenecks that influence the conservation of wheat landraces by farmers. This finding agrees with Awugachew Teshoma (2004) who indicated that the poor yield of the landraces and short rainy season period since most landraces are long maturing type would inhibit continuous production of the wheat landraces in some regions of Ethiopia. Claid and Ereck (2010) also explained that the expansion of improved bread wheat varieties and low soil fertility would inhibit the continuous production of the wheat landraces in Ethiopia. In addition, Wheat is the most dependable crop for the resource-poor highland farmers where poor soil fertility, frost, waterlogging, soil acidity, and soil degradation are the major yield-limiting factors, and where other cereals fail to grow (Woldeyesus Sinebo *et al*., 2010)

### 5.7. Naming of Wheat Landraces

Vernacular or local names were given for wheat landraces by farmers and have their meanings. For instance, ‘Qamadiguracha’ (Afaan Oromo) is named after its black seed color, ‘Inkoye’ is named after its very small seed, usually has good food quality. ‘Kenya’ most probably this variety is named after its original seed source. The names are simple and easily understood by farmers and are also important to maintain the identity of varieties. Negash Gelata and Grausgruber (2013) indicated that the wheat landraces could be named based on their seed colors, food quality, appearances of the head, and local name.

## 6.1. Conclusion

Digalu and Tijo is the district with diverse wheat landraces cultivation area of Arsi Zone, Ethiopia. Of these, wheat landraces growing in *Dega* kebeles were found to be more diverse with an overall H’ value (1.34) than those collected from *Woynadega* kebeles with H’ value (1.09). Of 27 wheat landraces identified by farmers, only 9 landraces are cultivated presently. Among the available wheat landraces, ‘Qamadiguracha’ is predominantly grown in the district due to its high end-use qualities. The overall genetic erosion in wheat landraces reached 66.6% in Digalu and Tijo districts. The loss of wheat landraces was accounted to major factors: introduction of modern wheat varieties; the introduction of other more productive crops; landraces low productivity; and infertility of soil. Food quality, Pests (for example, birds and insects), diseases resistance, market value, and straw quality were factors that initiated the farmers to maintain the diversity of wheat landraces. However, the Preservation of these landraces is influenced by bottlenecks like seed selection systems, soil infertility, insufficient crop yields, and short rainy seasons.

## 6.2. Recommendation

The regeneration of soil fertility alongside the re-introduction of lost landraces; improvement of landraces, on-farm conservation(*in-situ*) by re-sowing, saving of seeds for the future, seed bank conservation(*ex-situ*) are suggested for the restoration of wheat landraces diversity in DigaluTijo District.

